# Attention or prediction? Characterizing the top-down influence of predictive context on speech encoding

**DOI:** 10.64898/2026.08.11.742969

**Authors:** Alyssa Horng, Wei-Ching Lin, Ilaria Benciolini, Jin Dou, Aaron Nidiffer, Edmund C. Lalor

**Affiliations:** Department of Brain and Cognitive Sciences, University of Rochester, Rochester, NY 14627, USA; Department of Mechanical Engineering, University of Rochester, Rochester, NY 14627, USA; Del Monte Institute for Neuroscience, University of Rochester, Rochester, NY 14627, USA; Department of Biomedical Engineering, University of Rochester, Rochester, NY 14627, USA; Department of Neuroscience, University of Rochester, Rochester, NY 14627, USA; Center for Visual Science, University of Rochester, Rochester, NY 14627, USA

## Abstract

Theories of predictive coding propose that perception is the process of inferring the causes of our sensory input by comparing that input with predictions derived from our internal models of the world. Such predictive processes are thought to play a central role in language comprehension, however, robust neurophysiological evidence for such processes, particularly during natural speech perception, remains limited. Previous work has suggested that the early auditory encoding of words in natural speech is influenced by their preceding linguistic context. However, it remains unclear whether this effect is driven by prediction *per se* or dynamic modulations of attention based on contextual uncertainty. To distinguish between these alternatives, we recorded electroencephalography from 17 healthy adults while they listened to slightly changed audiobook. Specifically, we identified and replaced several unsurprising content words with more surprising words. We quantified the early auditory encoding of words using speech-envelope reconstruction accuracy within 100-ms time window after word onset and examined its relationship to word surprisal and contextual uncertainty. We found that more surprising words showed enhanced early auditory encoding despite matched contextual constraint. Moreover, the temporal profile of this enhancement depended on when the incoming speech signal diverged from the predicted phonological sequence, consistent with the emergence of prediction-error responses. Linear mixed-effects modeling further revealed that word surprisal had a substantially stronger influence on early auditory encoding than contextual uncertainty. Together, these findings indicate that the early auditory encoding of words during naturalistic speech perception is more strongly associated with predictive computations than with uncertainty-driven attentional gain.

**Significance Statement:** During natural speech comprehension, contextual information influences how the brain processes incoming sensory input. However, whether this context-dependent modulation of the early auditory encoding of words reflects predictive computations or dynamic changes in attentional gain has remained unresolved. By combining a naturalistic speech paradigm with a stimulus manipulation that varies word surprisal while controlling contextual uncertainty, we show that the early auditory encoding of words is driven by word surprisal under matched contextual constraint. Moreover, the temporal dynamics of this modulation closely follow the point at which the incoming speech signal departs from the predicted phonological sequence. These findings provide neurophysiological evidence that predictive computations contribute to the context-dependent modulation of early auditory processing during natural speech comprehension.

## Introduction

Speech is an important and complex signal that facilitates human communication. While the dominant view of speech comprehension involves serial and parallel processing (Hamilton et al. 2021), converging behavioral and neurophysiological evidence points to an overarching role for prediction in natural speech processing, such that responses to incoming speech sounds are influenced by the preceding linguistic context (Kuperberg and Jaeger 2016). However, despite substantial progress in the past few decades, how the brain transforms speech into meaning through predictive processes remains unclear.

One of the most influential theoretical frameworks of predictive perception is the predictive coding model (Clark 2013; Friston 2010). This model proposes that higher-levels in the cortical hierarchy generate predictions about lower-level sensory inputs through top-down connections. These predictions are then compared against incoming sensory signals, and the mismatch between the two, termed prediction error, is propagated up the hierarchy to update internal models of the world. Through iterative minimization of prediction error, perception is thought to emerge from the integration of prior expectations and sensory evidence.

Consistent with this framework, recent studies suggest that top-down predictions may affect speech processing at early stages of the auditory hierarchy (Sohoglu and Davis 2020; Donhauser and Baillet 2020; Broderick et al. 2019; Synigal et al. 2026). For example, Broderick and colleagues reported that during natural speech listening, the early auditory encoding of words is modulated by their semantic relationship to the preceding context. A more recent study used a similar framework to show that the cortical tracking of speech envelopes is enhanced for more contextually surprising words, suggesting that this enhancement may reflect prediction error computations during speech perception (Synigal et al. 2026).

It remains unclear whether stronger responses to more surprising words is a signature of prediction-error computations. Within a predictive-coding framework, such stronger tracking would be reflective of larger prediction errors generated by a mismatch between top-down expectations and bottom-up sensory input. However, an alternative potential explanation is that stronger tracking of surprising words arises from increases in attentional gain driven by contextual uncertainty. Previous work has shown that sensory processing is enhanced when contextual uncertainty regarding an upcoming word is high (Donhauser and Baillet 2020). According to this view, listeners might allocate additional attentional resources when the upcoming word is difficult to predict (or vice versa), therefore modulating the neural encoding of incoming acoustic information (Manker 2019).

These two accounts are grounded in two distinct contextual variables: lexical surprisal and cohort entropy. Lexical surprisal quantifies how unexpected a word is, while cohort entropy quantifies the level of contextual constraint placed on an upcoming word. For example, the sentence “The dentist told me to brush my…” provides strong contextual constraint and therefore has low entropy. The expected next word “teeth” has low surprisal, whereas an unexpected next word such as “car” has high surprisal. In contrast, the sentence “Last night in my dream I saw a…” allows many possible next words and therefore has high entropy. We used these two distinct contextual variables to index prediction error and uncertainty-modulated attention. Specifically, lexical surprisal was used as a proxy for the magnitude of prediction error, while cohort entropy was used as a measure of contextual uncertainty and its potential influence on attention allocation. The goal of the present study was to distinguish between prediction-error and attentional- gain accounts of the context-dependent modulation of the early auditory encoding of words. More specifically, we aimed to test the hypothesis that enhanced encoding reflects prediction-error computations beyond any effect that might derive from attentional gain induced by contextual uncertainty. To test this hypothesis, we causally manipulated lexical surprisal while controlling cohort entropy in natural continuous speech. Specifically, we recorded electroencephalography (EEG) from participants while they listened to an audiobook in which a subset of content words with low surprisal values were replaced with content words with high surprisal values. To test whether stronger encoding occurs at the point when the incoming speech signal deviates from the predicted phonological sequence, we created two types of word substitutions: one that differed from the original word at the initial phoneme and another that shared the same initial phoneme before differing later. The early auditory encoding of words was then compared across different conditions.

## Methods

### Participants

A total of 21 healthy adults (7 males, 18-35 years old) participated in this study. Four participants were excluded due to technical issues during data collection, resulting in a dataset of 17 participants. Each participant provided written informed consent and reported having normal hearing, normal or corrected-to-normal vision, English as their first and main language, and no history of neurological disorders. Participants were compensated for their participation. All procedures were approved by the University of Rochester Research Subjects Review Board.

### Experimental design

Participants listened to ∼70 minutes of *A Wrinkle in Time* by Madeleine L’Engle which was read in the voice of an American female speaker by Google Cloud’s text-to-speech AI software (voice = en-US-Journey-O). The stimulus was presented across 70 trials, each ranging from 47 to 68 seconds in duration with each trial beginning where the previous trial ended. To dissociate prediction error from dynamic attention based on contextual uncertainty, we aimed to manipulate lexical surprisal while controlling contextual uncertainty (cohort entropy). To do this, we identified six target words for replacement in each trial by ranking content words by lexical surprisal and selecting the six lowest. If two consecutive words were chosen, the second was kept intact and the word for replacement was re-selected. The newly inserted replacement words were selected by hand to preserve syntactic flow with the criteria that the lexical surprisal of the new word was above the median but within two standard deviations of the mean of the overall surprisal distribution.

Of the six changed words, three were replaced with words sharing the same initial phoneme as the original (e.g., “bed” replaced by “beacon”), and three with words differing in initial phoneme (e.g., “night” replaced by “century”; see a full example trial in Supplementary Material S1). For our analysis, we also selected three additional low-surprisal words in each trial as controls. We therefore based our primary analyses on three categories of selected words: changed words with different initial phoneme (the “changed_diff.”), changed words with the same initial phoneme (the “changed_same”), and unchanged words (the “unchanged”). Each category consisted of 210 words across trials (i.e., an average of 3 “changed_diff.” words, 3 “changed_same” words, and 3 “unchanged” words per minute).

After each trial participants answered two multiple choice comprehension questions. The comprehension questions were adapted from those used in two previous studies (Maddox and Lee 2018; Synigal et al. 2026) with minor adjustments to align them with the content of the altered words. The stimuli were presented through Sennheiser HD650 headphones at a sampling rate of 44.1 kHz using Psychtoolbox (Kleiner et al. 2007) and custom MATLAB scripts (MATLAB 2024).

### Computing lexical surprisal and cohort entropy

Lexical surprisal is a quantity describing how ‘surprising’ a word is. It is defined as the negative logarithm of a word’s probability of occurrence.

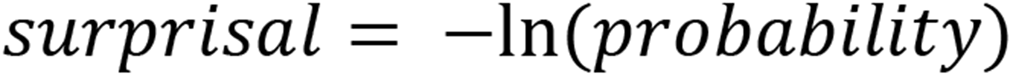

A word’s probability of occurrence given prior contextual information was estimated by using the large language model GPT-2 (Radford 2019). In practice, GPT-2 predicts the occurrence of a “token” instead of a word. A token is a common piece shared by words and is the minimal unit that GPT-2 used to model natural language. Approximately 30% of the words analyzed in this study were represented by two GPT tokens, whereas the remaining 70% were represented by a single GPT token. The lexical surprisal of the *j^th^* token is calculated as the conditional probability based on its preceding tokens.

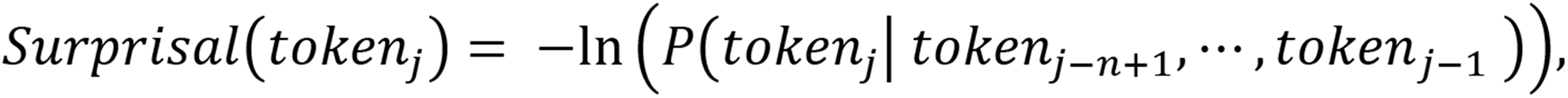

where *n* is the default maximum context window of GPT2 (*n =* 1024).

For a word having more than one token, we summed the surprisal values of its composing tokens to represent that word’s surprisal value. Overall, the lexical surprisal for the *i^th^* word is:

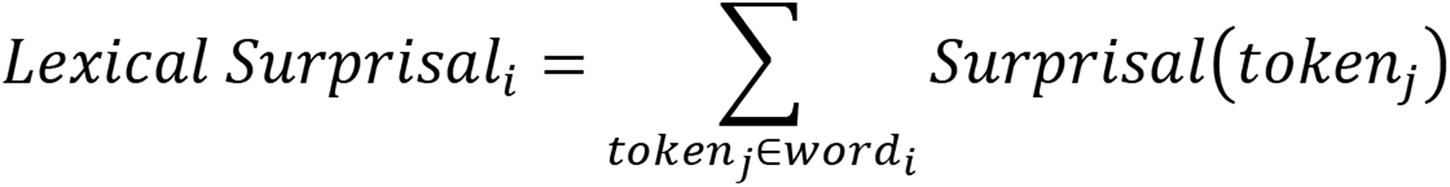

Besides lexical surprisal, to estimate how a word is contextually constrained given its preceding context, the cohort entropy of each word was also calculated. Cohort entropy describes the uncertainty that arises from the number of words compatible with the current context. For example, the context “The dentist told me to brush my…” has very low cohort entropy as there are very few words in real life that are compatible with that context. Meanwhile, a phrase like “Last night I had a dream and in my dream I saw a…” has very high entropy. The cohort entropy of words in our stimuli was calculated with GPT-2 using context-based predictability based on the first token of each word. The cohort entropy of the *i^th^* word is:

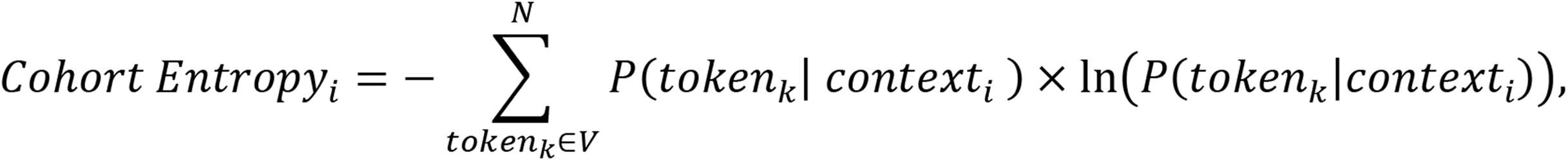

where *N* indicates the size of the cohort, *V* indicates GPT2’s token dictionary, *token_k_* indicates the *k^th^*token in the dictionary, and *context_i_* indicates the *i^th^* word’s current context which is equal to all preceding tokens within the maximum context window of GPT2.

### Data acquisition and preprocessing

EEG data were recorded from 128 scalp electrodes in addition to two mastoid channels used for re-referencing. The data were acquired at a 512 Hz sampling rate with the BioSemi Active Two system. An Arduino Uno microcontroller sampled the headphone output from the PC to detect an audio click at the start of each soundtrack to synchronize the BioSemi system with the start of each trial.

The data were preprocessed using functions from EEGLAB’s toolbox in MATLAB (Delorme and Makeig 2004). Firstly, the EEG was band-pass filtered using a fourth order Butterworth filter (1–8 Hz). It was then epoched to align with the 70 trials of audio and down sampled to 128 Hz. Finally, channels with abnormal (exceeding 5 standard deviations from the mean) kurtosis, spectral power, and amplitude were removed and recreated using spherical spline interpolation.

### Time-locked EEG (event-related potential) analysis

To validate that participants were attending to the audiobook narrative and finding the changed words surprising, we first carried out an event-related potential (ERP) analysis. In this analysis, EEG data was segmented into epochs time-locked to the onset of each word. The onset of each word was extracted from the speech stimulus using the Montreal Forced Aligner (McAuliffe et al. 2017) and inspected manually in Praat (Boersma and Weenink 2003). Epochs of the EEG data spanned 100 ms pre-onset and 800 ms post-onset, and an epoch was rejected for a given EEG channel if it contained EEG values outside the range -100 to 100 µV. ERPs were computed separately for each category of selected words (the “unchanged”, “changed_same”, and “changed_diff.”). For each participant, ERPs were obtained by averaging across 210 epochs at each time point within a category and were then averaged across participants.

### Defining an index of low-level auditory encoding at the word level

In order to quantify the degree of auditory encoding at the word level, we followed the methodology applied by previous studies (Broderick et al. 2019; Synigal et al. 2026). To evaluate neural responses to speech, we first quantified the speech in terms of its envelope. The envelope is a speech feature derived from the acoustic speech signal and commonly used to measure cortical activity related to speech processing (Aiken and Picton 2008; Destoky et al. 2020; Di Liberto et al. 2015; Ding and Simon 2013; Etard and Reichenbach 2019; Lalor and Foxe 2010; Nourski et al. 2009; Pasley et al. 2012). We calculated the envelope by resampling the speech signal to 128 Hz and averaging the square of the nearest neighbors every 4 samples, taking the square root and logarithmically scaling the RMS intensity (Lalor and Foxe 2010).

We trained a decoder, or “backwards” temporal response function (TRF) model, to reconstruct the speech envelope from the associated EEG data:

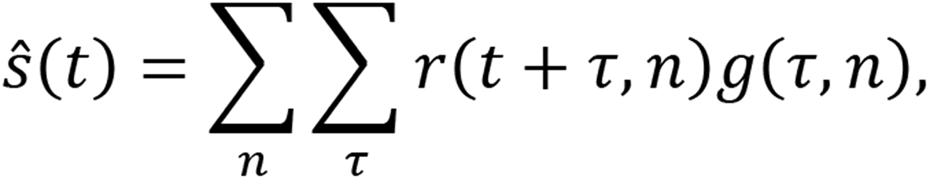

where *^S*(*t*) is the reconstructed estimate of the speech envelope, *g*(*τ*, *n*) is the TRF decoder, which represents the linear mapping from the lagged EEG responses *r*(*t* + *τ*, *n*), back to the stimulus. Here, the decoder integrates the neural response over a specified range of time lags *τ* and across the 128 EEG channels (Crosse et al. 2016).

The TRF model was trained and tested using a leave-one-out cross-validation procedure, in which the model trained on a rotating combination of 69 of the trials and was tested on its ability to reconstruct a speech envelope from the left-out trial. The EEG was lagged from 0 to 400 ms. A critical factor when training models in this way is to regularize those models so that they don’t overfit to the training data and, thus, generalize better in predicting unseen data. During training, we regularized our models using ridge regression, selecting an optimal regularization parameter (often denoted λ) from the range 10^0^–10^6^ using nested cross-validation, with the selected value for each participant maximizing the mean envelope reconstruction accuracy across channels and trials (see Crosse et al., 2016, 2019 for details). For the purposes of optimizing λ, model performance was quantified using reconstruction accuracy, defined as the Pearson correlation between the reconstructed and original speech envelope for each trial. The above procedures were implemented in MATLAB using the mTRF toolbox (Crosse et al. 2016).

After reconstructing the speech envelope for each trial, we defined a word-level index of early auditory encoding, termed “encoding index (EI),” based on word-level speech-envelope reconstruction accuracy within the first 100 ms after word onset. This time window was chosen to capture the fast-changing dynamics of early auditory encoding before recognition and resulting attentional changes (Broderick et al. 2019; Heinks-Maldonado et al. 2006) and was reported to show an apparent neural signature of prediction error computations (Sohoglu and Davis 2020). To investigate whether EEG is also sensitive to prediction errors that emerge later within a word, we additionally examined the index in the 100–200 ms post-onset interval. This analysis was motivated by the possibility that prediction-error responses may occur at different latencies depending on when an unexpected word diverges from the predicted phonological form.

Unlike previous studies (Broderick et al. 2019; Synigal et al. 2026), which quantify the word-level speech-envelope reconstruction accuracy using Spearman correlation between reconstructed and original envelopes, we instead employed the normalized mean absolute error (NMAE). NMAE is defined by the following equation:

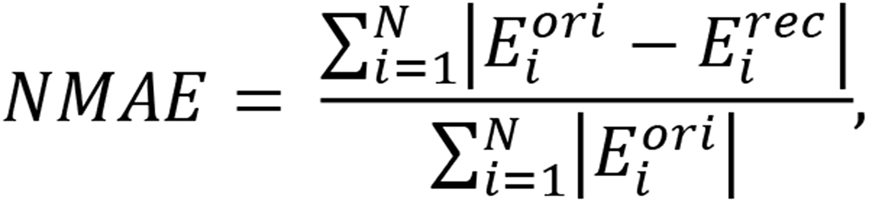

where *N* indicates the number of samples in the 100-ms time window, 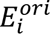 indicates the *i^th^*sample in the time window from the original envelope, while 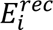 indicates the *i^th^* sample in the time window from the reconstructed envelope. Normalization (dividing mean absolute error by 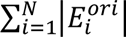) makes the quantity comparable across words with different envelope magnitude. Because larger NMAE values indicate poorer reconstruction, we defined word-level post-onset reconstruction accuracy—and thus the EI—as the negative NMAE:

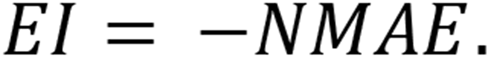

The original and reconstructed envelopes were z-scored and band-pass filtered (1–8 Hz) at the trial level prior to the analysis to ensure a fair comparison between the two signals. We adopted NMAE in place of Spearman correlation because the short analysis window (100 ms; ∼13 samples) renders correlation estimates unstable. In contrast, NMAE provides a more robust measure of the absolute deviation between reconstructed and original envelopes under these conditions. See Supplementary Material (S2) for further discussion.

### Relating early auditory encoding of words to their lexical surprisal

After defining the encoding index (EI) as the negative NMAE, we examined the relationship between EI and lexical surprisal using Pearson correlation. A significant positive correlation would indicate that words with higher lexical surprisal elicit larger encoding indices, consistent with findings from previous studies (Synigal et al. 2026).

### Statistical analysis

We confirmed that the changed words were as similarly constrained by context as the unchanged words by using independent-samples t-tests to compare cohort entropy across the word categories (the “unchanged”, “changed_same”, and “changed_diff.”). No significant differences in cohort entropy were observed between any pair of categories (“unchanged” vs “changed_diff.”: p = 0.55; “unchanged” vs “changed_same”: p = 0.38; “changed_diff.” vs “changed_same”: p = 0.75). See Supplementary Material (S3) for further details.

To test whether participants’ performance on the comprehension questions was above chance (25%), response accuracy was compared against chance level using a one-sample t-test across participants.

To assess whether the speech envelope reconstruction accuracy exceeded chance, we computed permutation-based null distribution of reconstruction accuracies for each participant by misaligning the reconstructed and original speech envelopes across trials. The mean of this null distribution was taken as the participant’s chance reconstruction accuracy. Group-level significance was then assessed by comparing the observed and chance reconstruction accuracies across participants using a pairwise t-test.

To examine whether lower-level auditory encoding differed across word categories (the “unchanged”, “changed_same”, and “changed_diff.”) and post-onset intervals (0–100 ms and 100– 200 ms), median EI was computed for each participant and compared using pairwise t-tests across the six resulting conditions (3 categories × 2 time windows).

Because the above analysis focused only on the “unchanged”, “changed_same”, and “changed_diff.” words, it did not allow us to test how EI might be affected by cohort entropy, which was constant across these words by design. To tease apart how prediction and attention might both affect the early auditory encoding of words, we conducted a complementary analysis using linear mixed-effects (LME) modeling across all the content words in our stimuli. This approach allows us to estimate the relationship between the main variable(s) of interest while accounting for variability due to random factors. In this study, the dependent variable was EI, defined as the negative NMAE between the reconstructed and original speech envelopes within the first 100 ms after word onset. The fixed effects included lexical surprisal and cohort entropy. To account for between-participant variability, the model also included by-participant random intercepts. In practice, the model was implemented using the fitlme function in Matlab, with the following equation:

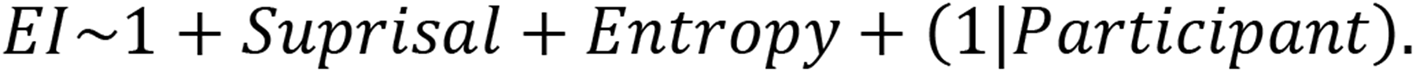

The dependent variable and predictors were normalized to have unit variance prior to the analysis. And, as mentioned above, the analysis was restricted to content words.

All the above-mentioned statistical analyses were performed in MATLAB R2024b (MATLAB 2024).

## Results

The present study aimed to test the hypothesis that enhanced early auditory encoding of words reflects prediction-error computations beyond any effect that might derive from attentional gain modulated by contextual uncertainty. We test this hypothesis by varying lexical surprisal while controlling contextual constraint. To be specific, participants listened to an audiobook while EEG was recorded. A subset of low-surprisal content words was replaced with high-surprisal content words while preserving the contextual constraint of the preceding context. Two types of word substitutions were created: one that differed from the original word at the initial phoneme (the “changed_diff.”) and another that shared the same initial phoneme before differing later (the “changed_same”). Speech envelopes were reconstructed from the EEG data using a backward TRF model. Early auditory encoding was quantified as the negative normalized mean absolute error (NMAE) between the original and reconstructed speech envelopes within post-onset time windows. The encoding index then was compared across different conditions.

### Participants comprehended the narrative and captured the substituted high-surprisal words

On average, participants answered 79.4% of the 140 comprehension questions correctly, which was significantly above the 25% chance baseline (one-sample t-test, p = 3.64×10^-11^). This indicates that participants attended to and comprehended the narrative content throughout the experiment, and that the narrative of the story was not destroyed by modifying a small percentage of the words.

To assess whether participants detected the semantic incongruities introduced by the changed high-surprisal words, an event-related potential (ERP) analysis was conducted. Semantically unexpected words are known to elicit an N400 response, characterized by a centro-parietal negativity approximately 400 ms after word onset (Frank et al. 2015; Kutas and Federmeier 2011; Kutas and Hillyard 1984). Accordingly, we examined ERPs at the midline parietal scalp (equivalent to channel Pz, where the N400 is typically strongest) by averaging EEG responses over all words and all participants for each substitution condition.

We observed clear neurophysiological signatures of semantic processing through an N400 response in the EEG data. ERPs elicited by changed words (both the “changed_diff.” and “changed_same” categories) showed pronounced negativities around 250–600 ms post-onset, while ERPs elicited by unchanged words showed no such negativity during this interval. In addition, the negativity was larger in amplitude and started to decline slightly earlier for the “changed_diff.” category than for the “changed_same” category (Figure 2A). Topographic maps further showed a centro-parietal negativity during the interval of 425-475 ms. Specifically, waveforms obtained by subtracting ERPs to unchanged words from ERPs to changed words display a scalp distribution characteristic of the N400 component (Figure 2B). Together, the behavioral and electrophysiological results indicate that participants successfully comprehended the broader narrative context while capturing the local semantic violations introduced by the stimulus manipulation.

**Figure 1.**
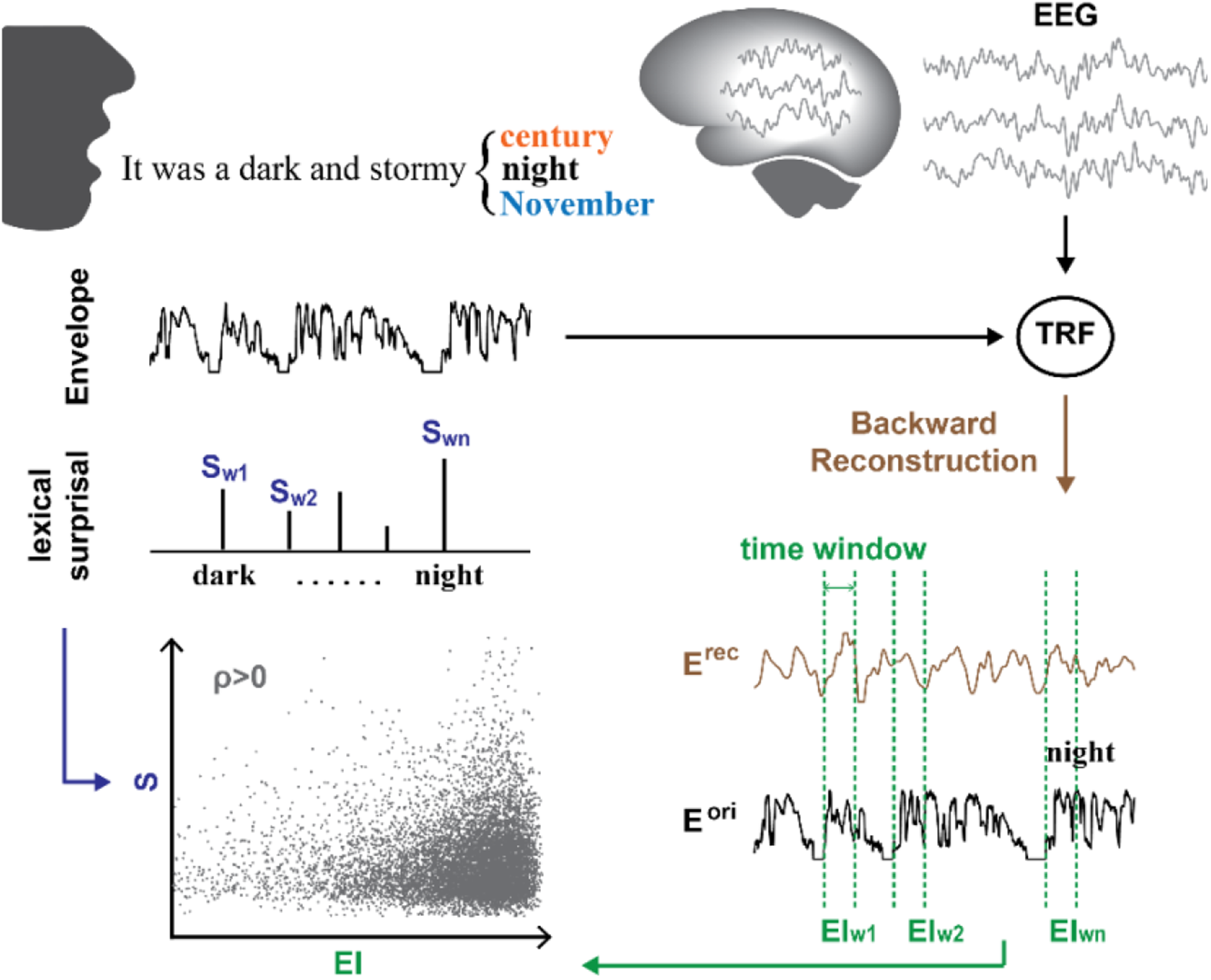
Experimental paradigm. EEG data were recorded while participants listened to an audiobook changed to have low-surprisal words (black) replaced with high-surprisal words (orange: with different initial phoneme; blue: with the same initial phoneme). A backward TRF model was used to reconstruct the speech envelope of the stimulus from the EEG data. Encoding index (EI, assessed by calculating the normalized mean absolute error between original and reconstructed speech envelope) was used as an early auditory encoding metric. The correlation between the EI and the context-based lexical surprisal of each word (calculated by GPT-2) was computed to determine the influence of lexical surprisal on early auditory encoding.

**Figure 2.**
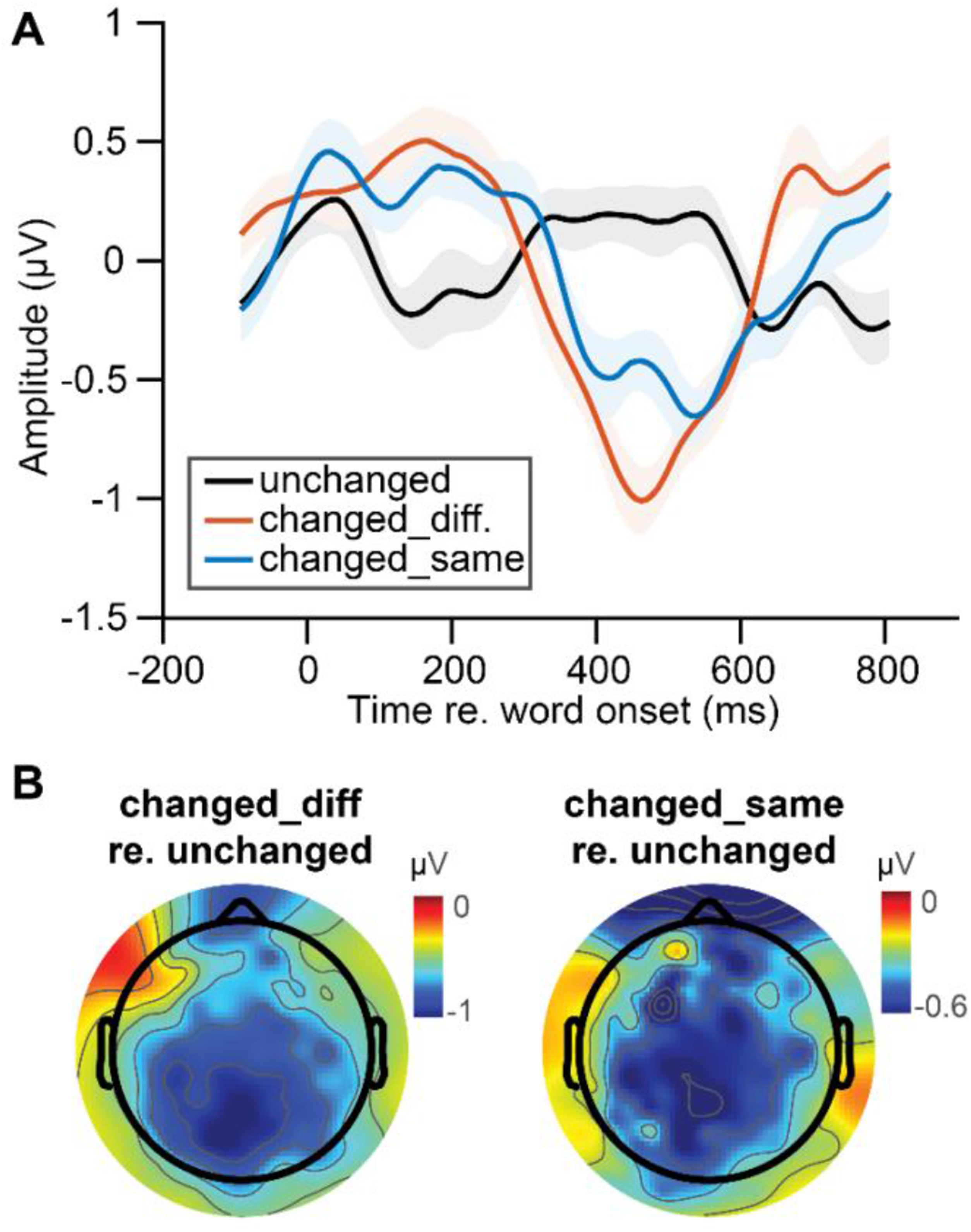
ERP analysis. (A) Grand average waveforms from channel Pz (center–back of the scalp) for the changed high-surprisal words (the “changed_diff.” and “changed_same”) and the unchanged low-surprisal controls (the “unchanged”). The classic N400 response was shown in both groups of changed words. Shaded area represented standard error for waveforms across participants. (B) Topographic maps of the N400 component (subtracting ERPs to unchanged words from ERPs to changed words) averaged over all trials and all participants over the interval 425–475 ms.

### Cortical tracking of the speech envelope is significantly above chance, while early auditory encoding of words increases as their lexical surprisal increases

Our primary test involves relating the effect of semantic context on the processing of the speech envelope across several conditions. As a prerequisite, we wished to verify that the EEG data exhibited reliable cortical tracking of the speech envelope. We thus first reconstructed the speech envelope from the recorded neural responses using a backward TRF model. Reconstruction accuracy at the trial level was quantified as the Pearson correlation between the reconstructed and original speech envelopes. We found that the reconstructed envelopes were positively correlated with the original speech envelopes for all participants (Figure 3A). Moreover, the distribution of trial-level reconstruction accuracies was significantly greater than chance (pairwise t-test, p = 8.48×10^-12^), indicating that EEG signals reliably tracked the temporal dynamics of the speech envelope. These results replicate previous findings of robust cortical tracking of natural speech (Aiken and Picton 2008; Destoky et al. 2020; Di Liberto et al. 2015; Ding and Simon 2013; Etard and Reichenbach 2019; Lalor and Foxe 2010; Nourski et al. 2009; Pasley et al. 2012) and validate the use of envelope reconstruction as an index of low-level auditory encoding in the present dataset. The central goal of our study was to further understand the mechanisms underlying previous published findings regarding the influence of predictive context on speech encoding. Specifically, previous work has shown that the cortical tracking of the acoustic envelope was stronger for words with higher lexical surprisal (Synigal et al. 2026). Therefore, before tackling our main question, we wished to verify that more surprising words resulted in stronger tracking of the speech envelope in our data. To investigate this, an encoding index (EI) was quantified by comparing the reconstructed and original speech envelopes within the first 100 ms after word onset. Specifically, EI was defined as the negative normalized mean absolute error (-NMAE) between the two envelopes. Then, for each participant, the relationship between EI and lexical surprisal was assessed using Pearson correlation. The analysis revealed a significant positive correlation between EI and lexical surprisal for all 17 participants (r = 0.05 ± 0.004; all participants: p < 10^-5^, Figure 3B). These results indicate that more surprising words have more accurate reconstruction of their post-onset speech envelopes. These findings replicate previous evidence that indices of low-level speech tracking are enhanced for more contextually surprising words (Synigal et al. 2026).

**Figure 3.**
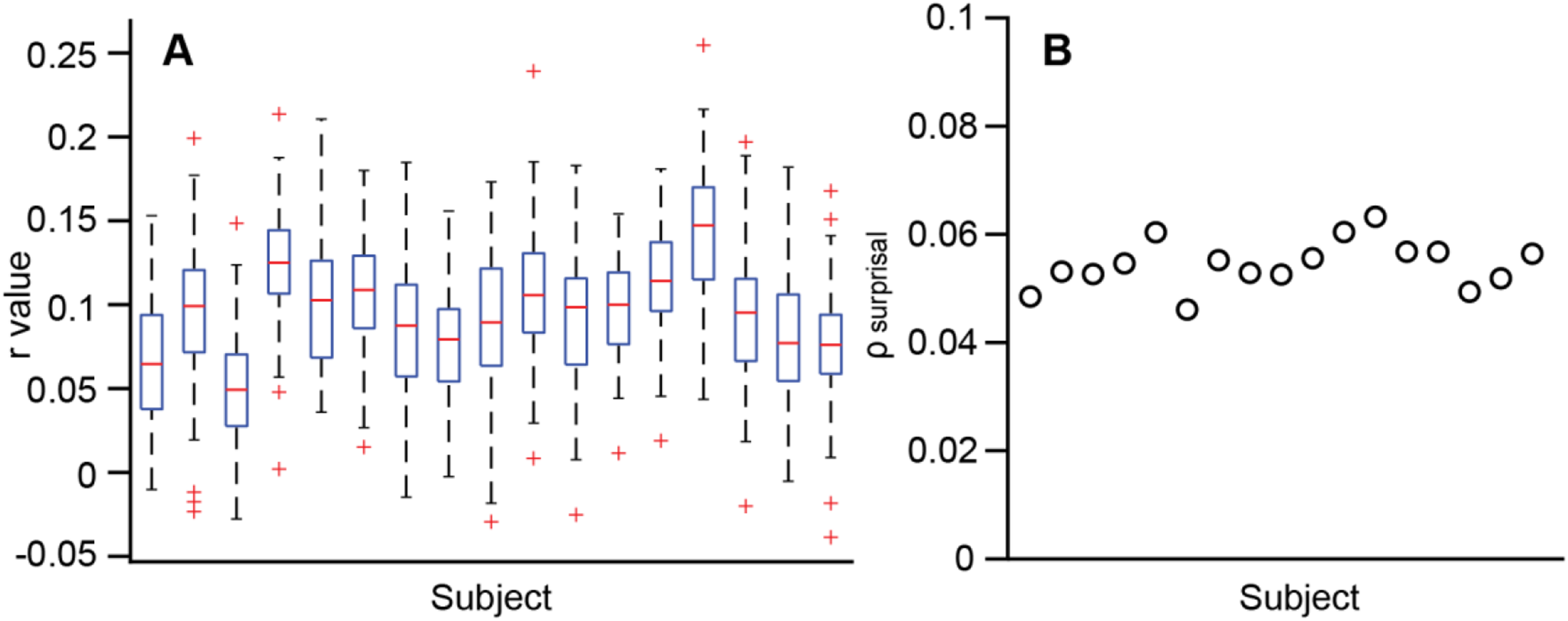
Cortical tracking of speech envelopes at the trial and the word level. (A) Box-and-whisker plots show trial-level reconstruction accuracy of speech envelopes reconstructed from neural responses for each participant. The red line indicates the median reconstruction accuracy, the box represents the interquartile range (25^th^-75^th^ percentiles), and the whiskers extend to the minimum and maximum non-outlying values across trials. Outliers are marked by red crosses. (B) Encoding index (EI) and lexical surprisal showed positive Pearson correlations (ρ_surprisal_) for all participants.

### Lexical surprisal modulates early auditory encoding under matched contextual constraint

To examine whether early auditory encoding differed across word categories (the “unchanged”, “changed_same”, and “changed_diff.”) and post-onset time windows (0–100 ms and 100–200ms), the median EI was computed for each participant and compared using pairwise t-tests across the six resulting conditions (3 categories × 2 time windows). The results are shown in Figure 4.

**Figure 4.**
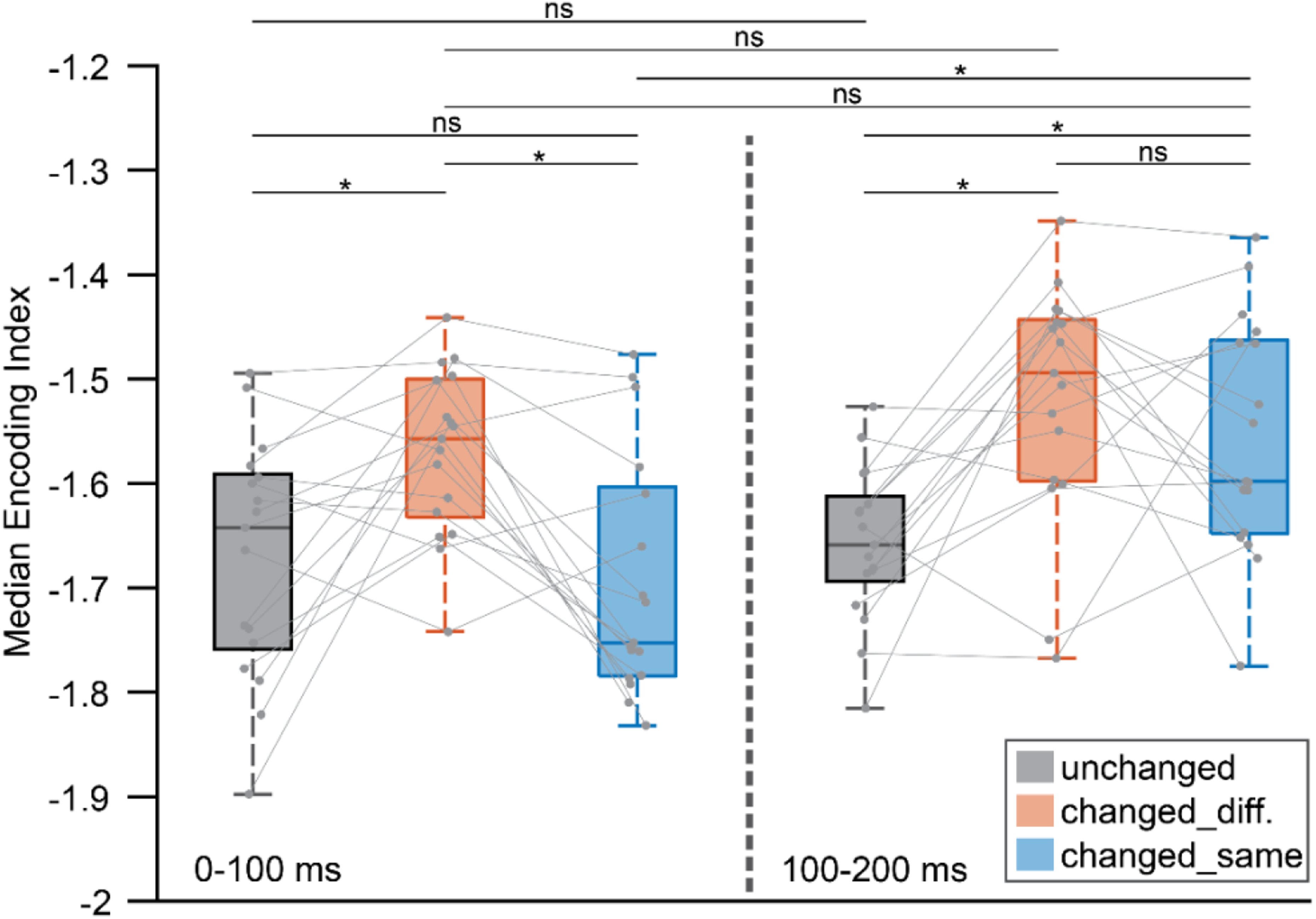
Participants’ median word-level EI for the six word-category × time-window conditions. Box-and-whisker plots show participant-specific median EI for the six conditions. The horizontal line within each box indicates the median across the 17 participants, the box represents the interquartile range (25^th^-75^th^ percentiles), and the whiskers extend to the minimum and maximum values across participants. Gray dots denote individual participants, and faint gray lines connect measurements from the same participant across conditions. Braces above the plots indicate the results of pairwise t-tests comparing median EI between conditions, with asterisks (*) denoting significant differences and “ns” indicating non-significant differences (p > 0.05).

To begin with, we examined whether words in the “changed_diff.” category elicited stronger EI immediately after word onset, where a prediction-error signal would be expected to emerge because the first phoneme already violates the predicted speech sequence. We found that within the 0–100 ms time window, words in the “changed_diff.” category had significantly greater median EI than words in the “unchanged” category (pairwise t-test, p = 0.0071). In contrast, no significant difference was observed between words in the “changed_same” and the “unchanged” categories during this interval (pairwise t-test, p = 0.51). These results suggest that the increase in EI depends on the timing of the phonological mismatch, supporting the interpretation that the observed effect reflects prediction-error computations.

Next, we examined whether words in the “changed_same” category elicited stronger EI later after word onset, as expected because the initial phoneme matches the listener’s expectation and the prediction is violated only once the speech diverges from the expected phonological sequence. We found that within the 100–200 ms time window, the “changed_same” category had significantly greater median EI than the “unchanged” category (pairwise t-test, p = 0.0086). Furthermore, median EI in the “changed_same” category was significantly greater in the 100–200 ms window than in the 0–100 ms window (pairwise t-test, p = 9.96×10^-5^). By contrast, neither the “unchanged” nor “changed_diff.” category showed a significant change in median EI between the two time windows (“unchanged”: p = 0.69; “changed_diff.”: p = 0.2). These results show that when the initial phoneme is consistent with the prediction, the enhancement is delayed until the mismatch occurs later in the word, consistent with the temporal dynamics expected from a prediction-error mechanism.

Finally, we compared the two conditions hypothesized to reflect prediction-error responses: the “changed_diff.” category in the 0–100 ms window and the “changed_same” category in the 100–200 ms window. We found that they did not differ significantly from one another (pairwise t-test, p = 0.63). This result implicates that prediction-error responses are comparable in magnitude once the incoming speech signal diverges from the predicted phonological sequence, regardless of when that divergence occurs.

Together, the above results suggest that lexical surprisal enhances indices of early auditory encoding while cohort entropy is controlled. Moreover, the temporal dissociation between the “changed_diff.” and “changed_same” conditions is consistent with the hypothesis that prediction-error signals emerge immediately when a word diverges from a predicted phonological form, as words sharing the same initial phoneme elicit stronger responses only after the prediction is violated. These findings therefore support the idea that the stronger tracking of the speech envelope for more surprising words reflects prediction-error.

### Early auditory encoding is more strongly influenced by lexical surprisal than cohort entropy

To assess the relative contributions of lexical surprisal and cohort entropy, we fit an LME model to the EI data across all content words in our stimuli. This approach allows the effects of lexical surprisal and cohort entropy on early auditory encoding to be estimated simultaneously while accounting for between-participant variability. Specifically, the model included by-participant random intercepts to capture differences in overall EI arising from participant-specific factors, such as anatomical variability, electrode sensitivity, and measurement noise. The results of the LME model are summarized in Table 1. To begin with, the model intercept was significantly lower than zero, indicating that EI remained negative after accounting for lexical surprisal and cohort entropy. Negative values of EI arise from its definition as the negative NMAE. Next, lexical surprisal showed a significant positive association with EI (β = 0.0374, p = 2.5×10^-27^), indicating that words with higher lexical surprisal tended to elicit stronger cortical tracking. This result is consistent with our analysis results that contextually surprising words evoke enhanced low-level auditory responses.

**Table 1.** LME fixed effects coefficients (95% CIs). LME model results show how the lexical surprisal and cohort entropy contribute to early auditory encoding of words.

|  | Estimate | Std. Error | t-Value | p-Value | Lower | Upper |
| --- | --- | --- | --- | --- | --- | --- |
| **Intercept** | -0.9555 | 0.0169 | -56.614 | 0 | -0.9886 | -0.9224 |
| **Surprisal** | 0.0374 | 0.0035 | 10.832 | 2.532e-27 | 0.0306 | 0.0442 |
| **Entropy** | -0.0075 | 0.0035 | -2.1583 | 0.031 | -0.0142 | -0.0007 |

In contrast, cohort entropy showed a significant negative association with EI (β = -0.0075, p = 0.031), indicating that words appearing in less constrained contexts tended to elicit weaker early auditory encoding, a result that speaks against the idea of a dynamic increase in attention for words that appear in unconstrained context. Additionally, the standardized effect size of lexical surprisal was approximately five times larger than that of cohort entropy. This suggests that even though both variables contribute to explaining variability in EI, lexical surprisal has a substantially stronger influence than cohort entropy.

## Discussion

The primary goal of this study was to determine whether the relationship between contextual predictability and early auditory speech encoding reflects prediction-error computations or whether it can be explained by uncertainty-driven attention gain. We hypothesized that prediction-error computations influence early auditory encoding beyond any influence of uncertainty-driven attentional gain. By manipulating lexical surprisal while controlling contextual constraint, we found that words with higher lexical surprisal showed stronger measures of cortical speech tracking despite matched cohort entropy. Furthermore, the temporal profile of this enhancement depended on the point at which the incoming speech signal diverged from the predicted phonological sequence. Specifically, stronger tracking emerged within the first 100 ms for words that differed from the expected word beginning at the initial phoneme, whereas it emerged later for words that shared the same initial phoneme. This temporal dissociation is difficult to explain with a generic attentional-gain account because the “changed_same” and “changed_diff.” categories (and, indeed, the “unchanged” words) were matched in contextual constraint and therefore should have had comparable levels of attention before word onset. The observed latency difference is consistent with a prediction-error account, in which enhanced auditory encoding arises only after the incoming speech signal provides evidence that violates the listener’s prediction. Together, these findings provide evidence that cortical tracking of words during natural speech perception is influenced by prediction-error computations.

Our experimental design varied lexical surprisal while controlling cohort entropy alone. As such, our results do not conclusively rule out a potential influence on EI from dynamic attention based on fluctuations in contextual constraint. One way to address this would be to carry out a similar study wherein one would vary cohort entropy while holding lexical surprisal constant. In the present study, we took a complementary approach, using an LME analysis to further investigate the relative contributions of lexical surprisal and cohort entropy across all content words in our audiobook stimuli. Importantly, the LME analysis revealed that lexical surprisal accounted for substantially more variance in EI than cohort entropy. This result suggests that prediction-error computations are more strongly associated with early auditory encoding than uncertainty-modulated attention. Moreover, the LME analysis revealed that the association between cohort entropy and EI was negative. This suggests that the lower the entropy the stronger the cortical tracking of the speech acoustics. Lower entropy implies higher constraint, meaning that the LME analysis suggests stronger EI for words occurring in more highly constrained contexts. This runs against our idea that lower constraints might lead to increases in attention, leading to stronger tracking of the speech (O’Sullivan et al. 2015). The fact that the contribution of cohort entropy to EI was significant at all indicates that attentional mechanisms may be playing a role in the early auditory processing of words in natural speech. However, we suggest that it is a task for future work to understand why this relationship is negative. Furthermore, it is also possible – even likely – that dynamic attention affects cortical tracking during natural speech processing in a way that does not relate to cohort entropy. Again, this is a question for future work.

The fact that lexical surprisal showed a much stronger association with early auditory encoding than cohort entropy is consistent with predictive-coding accounts in which prediction errors are suggested to be weighted by their precision. In this framework, perception is viewed as a Bayesian inference process where the prediction is the prior probability distribution, sensory input is the likelihood probability distribution, and the perceiver’s percept is the posterior probability (Sterzer et al. 2018). In the Bayesian scheme, prediction errors are suggested to be weighted by their precision (inverse variance of the probability distribution) which can either narrow their probability distributions (making them more precise) or widen the distributions (making them less precise). This means that the brain needs to encode not only the prediction errors themselves, but also the precision of these errors. Several researchers have proposed that attention serves as a precision weighting mechanism that can infer or estimate the certainty of the sensory evidence (Feldman and Friston 2010; Friston 2010; Hohwy 2012). From this perspective, attention and prediction error are not competing processes but complementary components of a unified inferential framework. Our LME results are broadly consistent with this view: although lexical surprisal had a substantially stronger influence on early auditory encoding than cohort entropy, cohort entropy also made a significant contribution. This finding suggests that early auditory encoding is influenced primarily by prediction-error computations while remaining sensitive to contextual uncertainty, potentially through attentional modulation of precision.

We confirmed that the replacement high surprisal words we inserted into the story were unpredicted by the participants by virtue of the fact that they produced an N400 response in our EEG data; topographic maps also displayed a scalp distribution characteristic (a centro-parietal negativity) of the N400 component. The presence of a robust N400 response provides independent evidence that participants captured the semantic incongruities introduced by the stimulus manipulation. This result supports the notion that N400 effects can occur during naturalistic speech listening without explicit tasks or listening instructions (Brodbeck et al. 2018), which alleviates the concerns that the N400 effect comes from “prediction-encouraging” experimental set-ups (Huettig and Mani 2016). On the other hand, the “changed_same” category had a smaller mean peak amplitude than the “changed_diff.” category, which is consistent with previous N400 study where Van Petten et al observed reductions in N400 responses to unexpected spoken words that shared the same initial phonemes as the expected completion (Van Petten et al. 1999).

The present study also makes methodological contributions. Previous work by Broderick et al developed a word-level measure of envelope reconstruction accuracy that enables the investigation of predictive processes on a word-by-word basis during naturalistic speech perception. We adopted a similar framework by using backward TRF models to reconstruct speech envelopes from the EEG recordings and then quantified early auditory encoding of words. However, unlike Broderick et al who quantified word-level reconstruction accuracy using Spearman correlation, we defined EI as the negative NMAE between the reconstructed and original speech envelopes. This modification provides a more robust measure of envelope similarity within short post-onset time windows, where correlation estimates can be unstable due to the limited number of samples. More broadly, the development of sensitive neurophysiological indices that can track online language processing during continuous, naturalistic speech remains an important goal in cognitive neuroscience. The present approach extends existing envelope- reconstruction methods by providing a more granular index of early auditory encoding that is sensitive to predictive processes during natural speech comprehension. This index may prove valuable for future investigations of predictive processing and could be particularly useful in terms of scalp EEG’s portable, non-invasive, and inexpensive nature.

In summary, the current results indicate that early auditory speech encoding during natural speech perception is more strongly influenced by prediction-error computations than by uncertainty-driven attentional gain. Predictive signals influence auditory processing at early stages of the speech-processing hierarchy and that this influence emerges at the point where incoming speech diverges from the listener’s expectations. Although contextual uncertainty also contributes to auditory encoding, its effect is negative and substantially smaller than that of lexical surprisal. Overall, these findings highlight the importance of distinguishing between top-down prediction and top-down attention and provide further evidence that prediction-error computations play a central role in shaping the neural processing of natural speech.

Future work could extend the results of the present study in several ways. First, as mentioned above, characterizing the role of cohort entropy more directly by manipulating contextual uncertainty while holding lexical surprisal constant would help to clarify the potential role of dynamic attention on early auditory encoding. In addition, investigating whether the influences of lexical surprisal and cohort entropy differ across neural frequency bands. For example, magnetoencephalography (MEG) studies have suggested that contextual uncertainty and surprise are associated with theta- and delta-band activity, respectively (Donhauser and Baillet 2020). Examining whether a similar dissociation is present in EEG measures of word-level auditory encoding may provide further insight into the neural mechanisms underlying prediction and attention during speech processing. Finally, investigating how these predictive effects operate across different levels of the speech-processing hierarchy. The present study focused on auditory encoding as indexed by speech-envelope reconstruction. Future work could extend this approach to higher-level representations, such as phonemic, lexical, or semantic processing, to better understand how predictive signals shape speech perception across multiple stages of cortical processing.

## Author contributions

A.H., I.B., A.R.N., E.C.L. designed research; A.H., W.C.L. performed research; W.C.L., A.H., J.D. analyzed data; W.C.L. created figures; A.H., W.C.L. wrote the first draft of the paper; A.H., W.C.L., A.N., E.C.L. edited the paper.

## Acknowledgements

This work was supported by a Schwartz Discover Grant from the University of Rochester and NIH R01 DC021140. The authors thank Ms. Sikhulile Vilane for assistance with data collection, Dr. Shyanthony Synigal, and Ms. Calli Smith for helpful guidance on data processing, and Dr. Samuel V. Norman-Haignere for helpful discussions on statistics.

## Supplementary Materials

### S1 Stimulus manipulation

Following paragraph is an example trial ∼1 min. The changed words are bold and color coded (“changed_same” in blue; “changed_diff.” in orange).

“It was a dark and stormy **century**. In her attic bedroom Margaret Murry, wrapped in an old patchwork quilt, sat on the foot of her **beacon** and watched the trees tossing in the frenzied lashing of the **whistle**. Behind the trees clouds scudded frantically across the sky. Every few **tomatoes** the moon ripped through them, creating wraithlike shadows that raced along the ground. The house shook. Wrapped in her **queen**, Meg shook. She wasn’t usually afraid of weather. It’s not just the weather, she thought. It’s the weather on top of everything else. On top of me. On top of Meg Murry doing everything wrong. School. School was all wrong. She’d been dropped down to the lowest section in her grade. That morning one of her **grocers** had said crossly, “Really, Meg, I don’t understand how a child with parents as brilliant as yours are supposed to be such a poor student.”

**Table S1.** Changed high-surprisal words in an example trial and their original words with lexical surprisal values.

| Original word | Original surprisal | Changed word | Changed surprisal |
| --- | --- | --- | --- |
| night | 1.77 | **century** | 12.06 |
| bed | 0.44 | **beacon** | 13.27 |
| wind | 2.04 | **whistle** | 12.23 |
| moments | 2.31 | **tomatoes** | 14.38 |
| quilt | 2.21 | **queen** | 15.41 |
| teachers | 1.21 | **grocers** | 15.56 |

### S2 Negative NMAE vs Spearman correlation as encoding index

We adopted negative NMAE in place of Spearman correlation because the short analysis window (100 ms; approximately 13 samples at the downsampled rate) renders correlation estimates unstable. In contrast, negative NMAE provides a more robust measure of the absolute deviation between the reconstructed and original speech envelopes under these conditions.

To illustrate this difference, Figure S1 shows the average standard error of participant-specific encoding index (EI) across participants as a function of the analysis-window length. When EI was quantified using Spearman’s correlation coefficient, the average standard error decreased steadily as the analysis window increased, indicating that the reliability of the correlation estimate was strongly dependent on the number of available samples (Figure S1A) across the range of time-windows. By contrast, when EI was defined as negative NMAE, the average standard error reached a stable plateau at approximately 250 ms, suggesting that negative NMAE provides a substantially more stable estimate of EI, particularly for short windows (Figure S1B).

**Figure S1.**
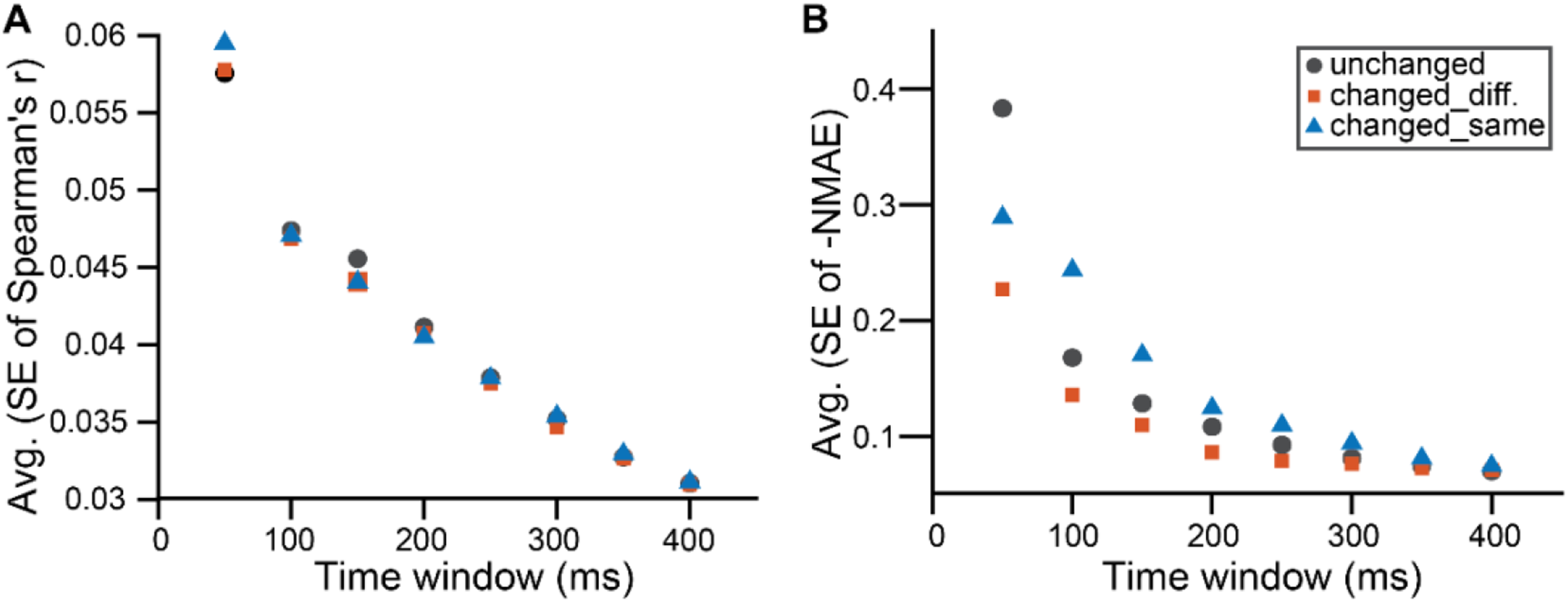
Average standard error of encoding index across participants for the selected word categories. using (A) Spearman correlation (B) negative NMAE.

### S3 Lexical surprisal and cohort entropy for selected word categories

To confirm that the changed words were matched to the unchanged words in contextual constraint, we compared cohort entropy across the three word categories (the “unchanged”, “changed_same”, and “changed_diff.”) using pairwise independent-samples t-tests. No significant differences in cohort entropy were observed between any pair of categories (“unchanged” vs “changed_diff.”: p = 0.55; “unchanged” vs “changed_same”: p = 0.38; “changed_diff.” vs “changed_same”: p = 0.75; Figure S2A), confirming that the changed and unchanged words were similarly constrained by their preceding context.

As expected, lexical surprisal differed between conditions. Both changed word categories (the “changed_same” and “changed_diff.”) exhibited significantly higher lexical surprisal than the unchanged words (“unchanged” vs “changed_diff.”: p = 2.8×10^-129^; “unchanged” vs “changed_same”: p = 2.06×10^-119^; Figure S2B), confirming that the stimulus manipulation successfully increased lexical surprisal while preserving contextual constraint. At the same time, for changed words, the “changed_same” exhibited significantly higher lexical surprisal than the “changed_diff.” (p = 1.83×10^-7^; Figure S2B).

**Figure S2.**
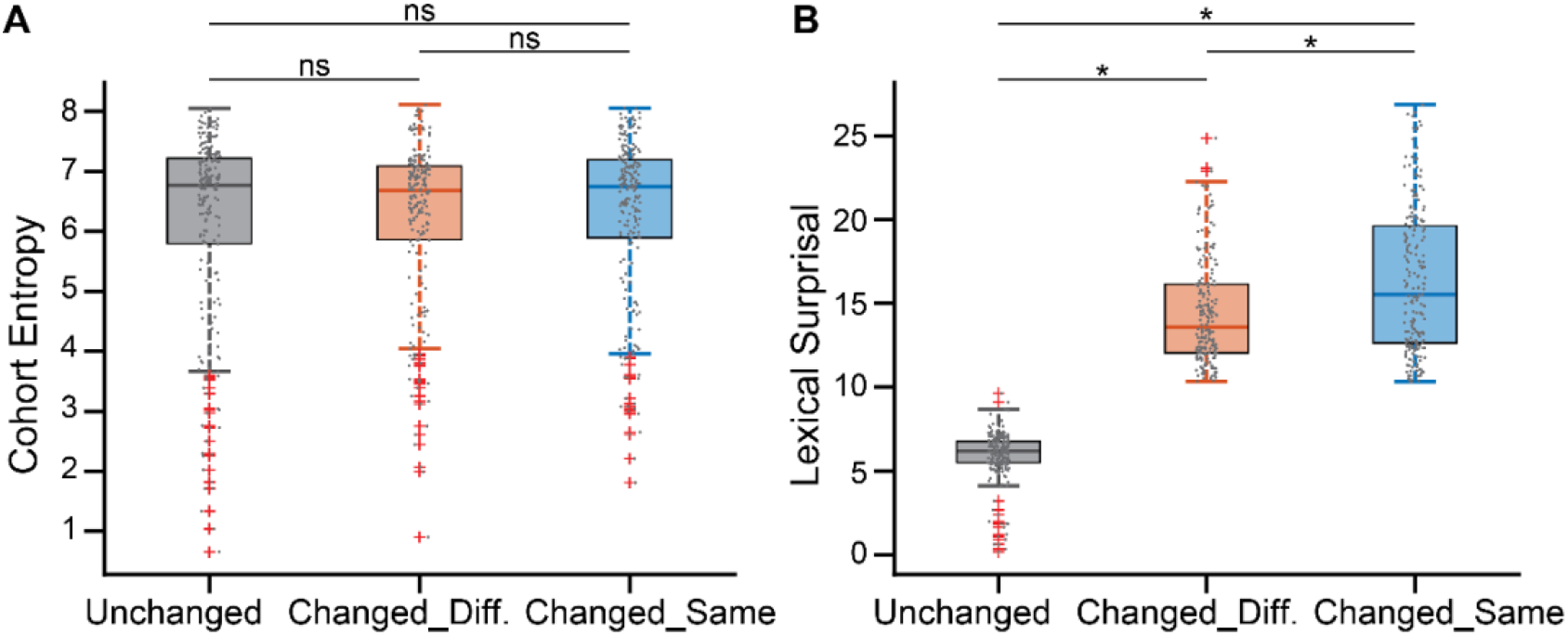
Cohort entropy and lexical surprisal of selected word categories.

